# Evaluation of N8-propargyl spermidine as a chemical probe for deoxyhypusine synthase-dependent post-translational modifications

**DOI:** 10.64898/2026.11.10.765603

**Authors:** Paweł Kochanowski, Marta Stolarek-Sipior, Magdalena Chojnacka, Przemysław Wanat, Przemysław Grudnik, Remigiusz Serwa

## Abstract

Hypusination is a highly conserved post-translational modification generated on eukaryotic translation factor 5A (eIF5A) by the sequential action of deoxyhypusine synthase (DHS) and deoxyhypusine hydroxylase (DOHH). Recently, expanded DHS substrate specificity and DHS-dependent acetylhypusination of receptor-interacting serine/threonine-protein kinase 1 (RIPK1) have been proposed based on studies employing N8-propargyl spermidine in click chemistry-based workflows, alongside additional analyses. Here, we used orthogonal approaches to evaluate N8-propargyl spermidine as a chemical probe for protein hypusination and to reassess the evidence for DHS-dependent modification of RIPK1. Protein labeling following treatment of cells with the probe was unaffected by DHS inhibition but was strongly reduced when cysteines were alkylated after metabolic labeling and before initiation of the click reaction, suggesting that probe-associated adducts form through reactions involving free cysteine thiols after cell lysis. Consistent with this observation, analysis of probe-modified peptides localized adducts to cysteine residues, with masses consistent with copper-dependent azide–alkyne–thiol coupling, and detected no lysine modifications corresponding to propargylated hypusine or deoxyhypusine. In a reconstituted enzymatic assay, N8-propargyl spermidine did not support DHS-dependent modification of eIF5A, whereas native spermidine supported robust modification of eIF5A but no detectable modification of RIPK1. Furthermore, our structural study revealed a non-productive binding mode of N8-propargyl spermidine to DHS. These findings challenge the interpretation of widespread chemical probe-derived labeling as evidence for expanded DHS substrate specificity and highlight the need for stringent validation before proposing new hypusination targets.

## Introduction

Hypusination is a unique and highly conserved post-translational modification (PTM) generated through sequential action of deoxyhypusine synthase (DHS)(1–4) and deoxyhypusine hydroxylase (DOHH)(1, 5, 6) on lysine 50 of eukaryotic initiation factor 5A (eIF5A)(7, 8). During this process, DHS transfers the 4-aminobutyl moiety from spermidine (SPD) to eIF5A to generate deoxyhypusine, which is subsequently hydroxylated by DOHH to form hypusine(9). Human DHS functions as a homotetrameric assembly with a dimer-of-dimers architecture and D2 symmetry. The active sites form at the dimer interfaces, with one subunit contributing most of the catalytic site and the other protomer providing the majority of the NAD-binding site(10). Extensive biochemical, structural, and genetic studies have established eIF5A as the canonical and apparently unique eukaryotic DHS substrate(11–16).

Tagging-via-substrate (TvS) is a chemoproteomic strategy that uses bioorthogonally tagged substrate analogs to study protein post-translational modifications(17). In copper-catalyzed azide– alkyne cycloaddition (CuAAC)(18, 19)-based TvS, an alkyne- or azide-bearing analog must remain compatible with the relevant enzymatic pathway and be metabolically incorporated into its protein targets. The introduced handle can then be conjugated to fluorescent or affinity tags for detection, enrichment, and proteome-wide identification by mass spectrometry. This strategy is particularly useful when alternative methods for enriching proteins carrying the modification of interest are unreliable, or when the dynamics of a reversible PTM need to be monitored, as demonstrated by us and others in studies of protein lipidation(20–25).

More recently, to facilitate the investigation of understudied aspects of polyamine metabolism, their interactions with proteins and subcellular trafficking, we developed a series of alkyne-tagged polyamine probes(26–28) compatible with CuAAC. One of these probes, N8-propargyl spermidine (Probe 1), which has also found application beyond polyamine biology(28), appeared particularly attractive for studying DHS-dependent PTM. If accepted as a substrate by DHS, Probe 1 could, in principle, enable transfer of a propargylated 4-aminobutyl moiety to eIF5A, thereby providing a potential chemical reporter of DHS-mediated modification.

Zhang et al. recently employed Probe 1 in a CuAAC-based proteomic workflow and proposed a substantial expansion of DHS substrate specificity, reporting hundreds of putative protein substrates, including receptor-interacting serine/threonine-protein kinase 1 (RIPK1)(29). As a key regulator of inflammation and cell death whose activity and protein interactions are controlled by PTMs(30, 31), RIPK1 represents an important target for identifying novel modifications and establishing their functional consequences(32, 33). Based on subsequent experiments without Probe 1, the authors proposed that Lys140 within the RIPK1 kinase domain carries a novel acetylated hypusine modification, which they termed “acetylhypusination”. They reported this modification to be spermidine-dependent, although the origin of the acetyl moiety was not established. The proposed modification was further suggested to inhibit RIPK1 activation and thereby reduce inflammatory and cell death-associated processes contributing to diabetes-associated pathology(29).

However, interpreting Probe 1-derived labeling as evidence of DHS-dependent modification requires confirmation that DHS can accept the analog and transfer its tagged moiety to a protein substrate. This cannot be assumed from structural similarity to spermidine alone, because DHS recognizes its natural substrate through an extensive network of interactions spanning nearly the entire molecule(9, 10, 13, 34). Equally important is establishing whether the detected labeling reflects this transfer reaction or arises through other mechanisms. We recently demonstrated Probe 1-derived covalent labeling of aldehyde dehydrogenase 1A1, showing that the labeling is not restricted to the proposed hypusination pathway(28). Additional labeling can arise during CuAAC itself, including through an azide–alkyne–thiol side reaction that generates cysteine-linked thiotriazole adducts under commonly used reaction conditions(35). Distinguishing these sources of labeling therefore requires chemical characterization of the probe-derived protein adducts to assess whether their masses match those predicted from the proposed tag-transfer mechanism. Such analysis is also valuable and often performed in TvS workflows because it can identify sites of probe incorporation.

These considerations prompted us to independently evaluate Probe 1 as a DHS substrate and reporter of DHS-dependent PTMs. Here, we combine chemical proteomics, biochemical reconstitution, and X-ray crystallography to investigate the molecular basis of probe-associated protein labeling and independently assess the proposed DHS-dependent modification of RIPK1.

## Results

### N8-propargyl spermidine promiscuously labels proteins during CuAAC independently of DHS activity

To evaluate the applicability of Probe 1 in cellular systems, we first examined Probe 1-derived protein labeling in mouse embryonic fibroblasts (MEFs), the principal cellular model used by Zhang et al.(29), and included HEK293T cells as an additional model. Following probe treatment, cells were lysed and subjected to CuAAC with azide-functionalized TAMRA or biotin for fluorescence detection or affinity enrichment, respectively. Numerous fluorescent protein bands were observed in both cell models (Fig. 1A–B), and complementary LC–MS/MS analysis identified hundreds of proteins enriched from MEFs following treatment with Probe 1 relative to untagged spermidine (Fig. 1C, Table S1), consistent with the broad enrichment previously reported in this model(29).

**Figure 1.**
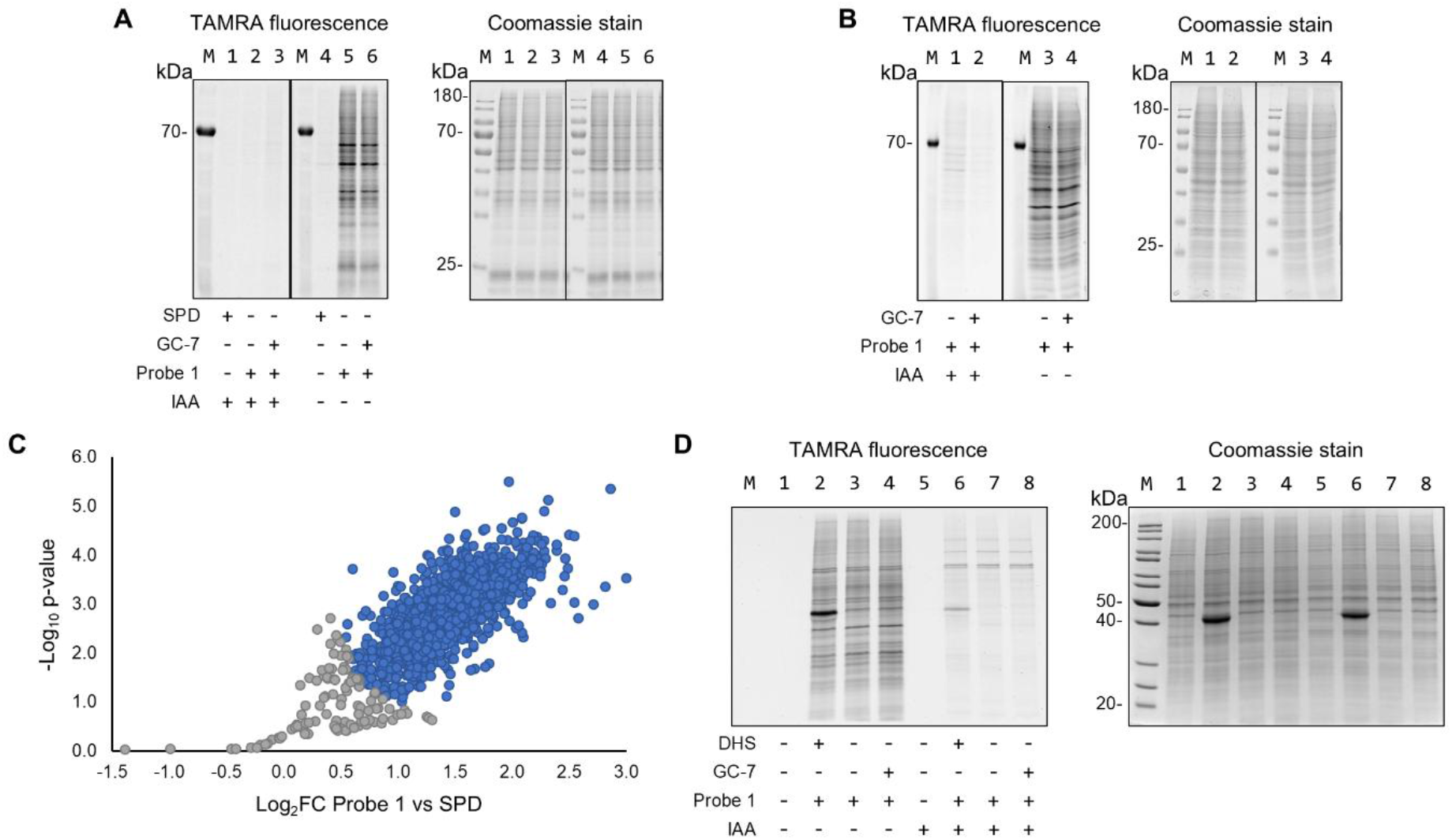
Cysteine capping-dependent formation of N8-propargyl spermidine-protein conjugates. **A.** TAMRA fluorescence after CuAAC ligation with lysate from Probe 1-treated MEF cells (left) and Coomassie brilliant blue staining of the samples (right). **B.** TAMRA fluorescence after CuAAC ligation with lysate from Probe 1-treated HEK293T cells (left) and Coomassie brilliant blue staining of the samples (right). **C.** LC-MS/MS analysis comparing protein level enrichment from MEF cells treated with Probe 1 or SPD after CuAAC ligation to biotin. Proteins significantly enriched in the Probe 1 condition (Student’s t-test, permutation-based FDR = 0.05, S0 = 0.5, n = 3) are shown in blue, other proteins are shown in grey. The numerical values used to generate the volcano plot are provided in Table S1. **D.** TAMRA fluorescence after CuAAC ligation with native lysate from HEK293T cells, incubated with Probe 1 and recombinant DHS (left), and Coomassie brilliant blue staining of the samples (right).

However, several observations suggested that a substantial fraction of these signals did not arise from DHS-mediated transfer reactions. Most notably, pretreatment of lysates with iodoacetamide prior to CuAAC strongly attenuated Probe 1 labeling across multiple experimental conditions (Fig.1A-B). This finding is consistent with our related work on propargylated polyamines, in which cysteine alkylation was deliberately omitted before CuAAC to enable LC-MS/MS analysis of probe-modified cysteine-containing peptides as readouts of potential intracellular interconversion of these compounds(27). In contrast, pretreatment with N1-guanyl-1,7-diaminoheptane (GC-7), a well-characterized DHS inhibitor(36), did not abolish Probe 1-derived signals (Fig.1A-B).

To further test whether DHS is required for Probe 1-protein conjugation, we incubated native HEK293T lysate with this compound. Under these conditions, similar extensive protein tagging was observed after the click reaction (Fig.1D). Addition of recombinant DHS resulted only in the appearance of a single additional fluorescent band at approximately 40 kDa corresponding to DHS monomer. Likewise, inhibition of endogenous DHS activity in the lysate by preincubation with GC-7 before Probe 1 addition did not affect the band pattern or band intensities (Fig.1D). In contrast, cysteine alkylation prior to click coupling with azide-functionalized TAMRA once again markedly reduced labeling in all samples (Fig.1D). Together, these findings indicate that the observed Probe 1 attachment to proteins arises primarily during CuAAC rather than from its enzymatic incorporation in cells.

### Proteomic analyses identify cysteine-associated N8-propargyl spermidine adducts after CuAAC

To assess the chemical identity of the observed modifications, we lysed Probe 1-treated HEK293T cells and subjected the lysate to CuAAC ligation with an azide-functionalized biotin tag, followed by tryptic digestion. Probe-modified peptides were then affinity-enriched and analyzed by LC– MS/MS. Database searches targeting biotin-tagged triazole adducts of propargylated deoxyhypusine or hypusine, the expected products of DHS-catalyzed transfer, as well as cysteine-linked thiotriazole adducts of Probe 1 identified 147 peptide sequences bearing cysteine-localized modifications with mass shifts consistent with the predicted thiotriazole adducts, whereas no peptides bearing the expected lysine-linked adducts were detected (Table S2). Copper-dependent azide–alkyne–thiol coupling has previously been described as a source of background labeling in CuAAC-based proteomics, and analogous adducts were detected in our earlier studies of propargylated polyamine probes(27, 35). Importantly, adduct formation strongly depended on assay conditions, as cysteine alkylation before CuAAC markedly reduced the detection of these modified peptides (Table S2). This dependence on free cysteine thiols, together with the identified adducts, supports CuAAC-mediated thiotriazole formation as a major source of the extensive protein labeling observed with Probe 1.

### DHS cannot use Probe 1 as a substrate or modify RIPK1

We next examined whether Probe 1 functions as a DHS substrate in a defined *in vitro* system containing recombinant human DHS, DOHH, and eIF5A - the only well-established DHS substrate. Following incubation of eIF5A with Probe 1 or SPD as a positive control under *in vitro* (deoxy)hypusination conditions, LC–MS/MS analysis showed that addition of DHS and DOHH markedly reduced the signal intensities of unmodified eIF5A peptides encompassing K50 in reactions containing SPD, consistent with efficient modification at this site. No corresponding decrease was observed with Probe 1, indicating no detectable conversion in this assay (Fig. 2A, Table S3). By comparison, the intensities of peptides not encompassing K50 remained largely unchanged, providing an internal control for the selective decrease in unmodified K50-containing peptides observed with SPD (Fig.2A). Moreover, in reactions containing SPD, DHS, and DOHH, eIF5A peptides encompassing K50 with mass shifts consistent with deoxyhypusination and hypusination were readily detected (Fig. 2B, Table S3). In contrast, no modifications consistent with transfer of the propargylated 4-aminobutyl moiety to eIF5A were detected in the corresponding reactions containing Probe 1. Together, these results suggest that, unlike the 4-aminobutyl moiety of spermidine, the propargylated 4-aminobutyl moiety of Probe 1 is not transferred to eIF5A under conditions optimized for DHS/DOHH-dependent modification.

**Figure 2.**
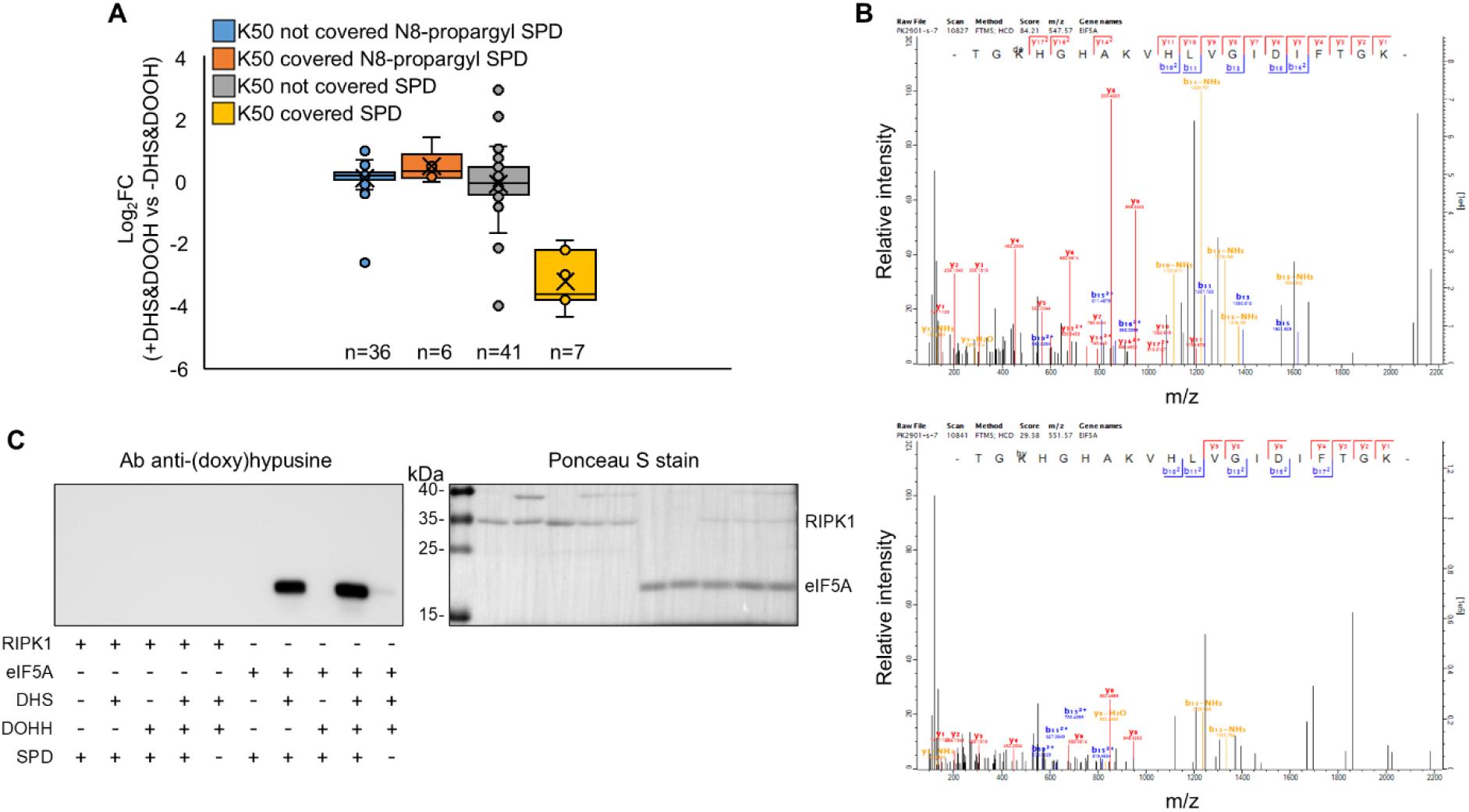
Probe 1 is not a DHS substrate, and RIPK1 does not undergo DHS/DOHH-mediated PTM. **A.** *In vitro* hypusination in the presence of spermidine, but not Probe 1, decreases the intensity of unmodified eIF5A peptides encompassing the known hypusination site in LC-MS/MS analysis. The distribution of peptide intensity ratios (complete reaction vs reaction without enzymes) was separated into two peptide populations: peptides encompassing the known hypusination site in eIF5A and peptides not encompassing this site. **B.** *In vitro* hypusination in the presence of spermidine, but not Probe 1, yields modified eIF5A peptides encompassing the known hypusination site. Known spermidine-derived modifications (hypusine and deoxyhypusine), as well as putative Probe 1-derived modifications (N-propargyl hypusine and N-propargyl deoxyhypusine), were searched for on eIF5A peptides. MS/MS spectra showing deoxyhypusination (upper panel) and hypusination (lower panel) on the eIF5A peptide encompassing the known modification site. No putative Probe 1-derived modifications were detected. **C.** Western blot-based *in vitro* (deoxy)hypusination assay using natural spermidine and a context-independent antibody that detects both deoxyhypusine and hypusine shows formation of (deoxy)hypusinated eIF5A in appropriate conditions, while no modification on RIPK1 can be detected.

Having confirmed that native SPD but not Probe 1 supports efficient eIF5A modification in the *in vitro* system described above, we next examined whether RIPK1 could also serve as a DHS substrate in reactions containing SPD. For that, we used western blotting with a context-independent antibody that recognizes both deoxyhypusine and hypusine regardless of the surrounding amino acid sequence(37). Although eIF5A (deoxy)hypusination was readily observed in parallel reactions, no modification of RIPK1 was detected (Fig. 2C). Thus, the assay supported modification of the established substrate but provided no evidence for DHS/DOHH-dependent modification of RIPK1 under the conditions tested.

### Non-productive binding mode of N8-propargyl spermidine to DHS

To understand N8-propargyl spermidine interaction with DHS at a molecular level, we co-crystallized this protein with its co-factor (NAD) and Probe 1. We collected diffraction data and solved the structure by molecular replacement using PDB 6XXJ as a search model. DHS crystallized in the P3_2_21 space group with one dimer per asymmetric unit and crystallographic symmetry generating the tetrameric biological assembly. The structure was refined to R_work_/R_free_ of 16.9%/19.7% at 1.61 Å (Table 1). The molecular model encompasses residues 28-363. Some signal for the preceding part of the N-terminal *ball-and-chain* motif is present in the map, but it does not allow for confident modeling of this region of DHS. The model contains two copies of Probe 1 bound symmetrically to the two active sites formed at the dimer interface

**Table 1.**
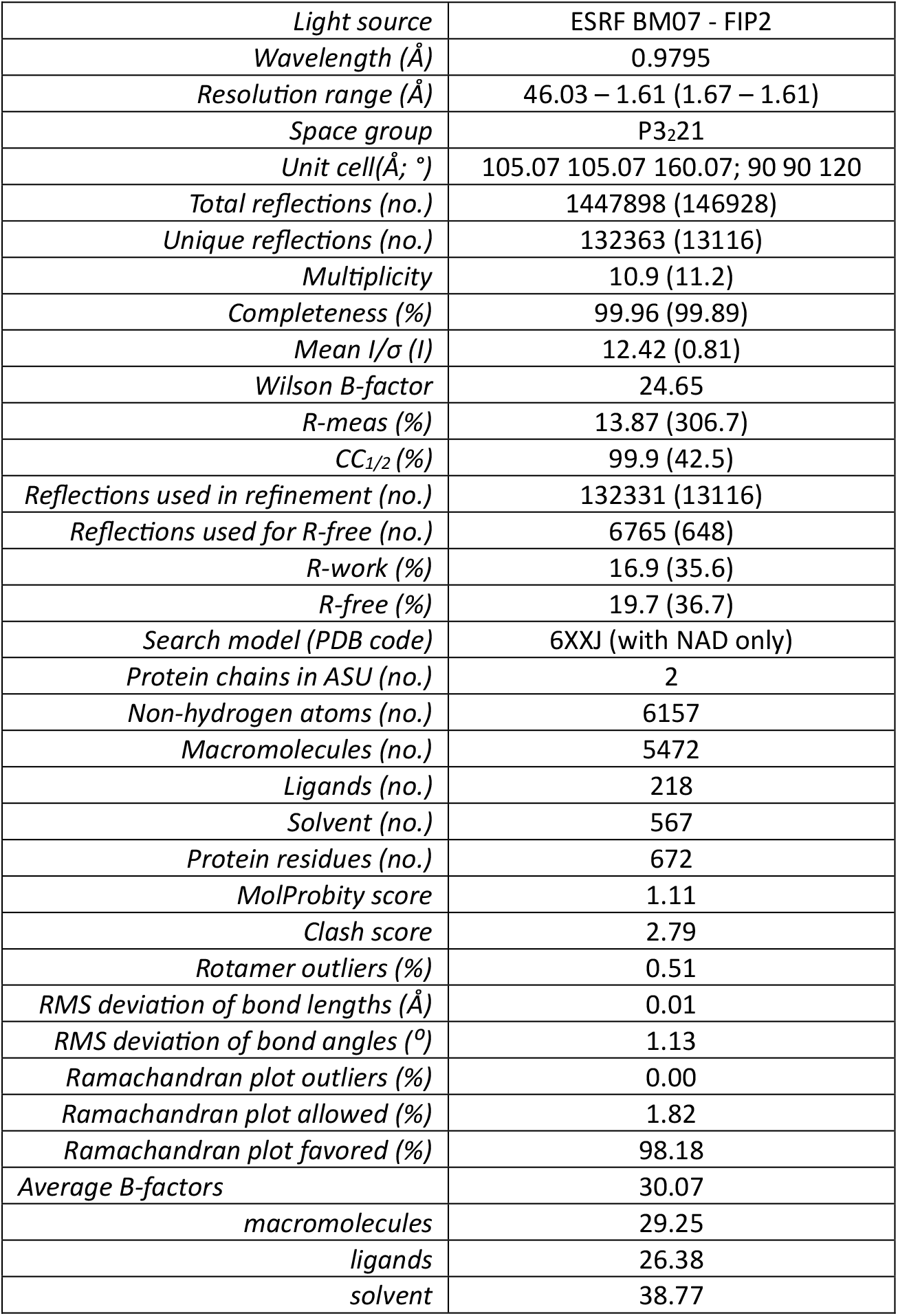
Statistics of the crystallographic data.

| *Light source* | ESRF BM07 - FIP2 |
|---|---|
| *Wavelength (Å)* | 0.9795 |
| *Resolution range (Å)* | 46.03 – 1.61 (1.67 – 1.61) |
| *Space group* | P3<sub>2</sub>21 |
| *Unit cell(Å; °)* | 105.07 105.07 160.07; 90 90 120 |
| *Total reflections (no.)* | 1447898 (146928) |
| *Unique reflections (no.)* | 132363 (13116) |
| *Multiplicity* | 10.9 (11.2) |
| *Completeness (%)* | 99.96 (99.89) |
| *Mean I/σ (I)* | 12.42 (0.81) |
| *Wilson B-factor* | 24.65 |
| *R-meas (%)* | 13.87 (306.7) |
| *CC<sub>1/2</sub> (%)* | 99.9 (42.5) |
| *Reflections used in refinement (no.)* | 132331 (13116) |
| *Reflections used for R-free (no.)* | 6765 (648) |
| *R-work (%)* | 16.9 (35.6) |
| *R-free (%)* | 19.7 (36.7) |
| *Search model (PDB code)* | 6XXJ (with NAD only) |
| *Protein chains in ASU (no.)* | 2 |
| *Non-hydrogen atoms (no.)* | 6157 |
| *Macromolecules (no.)* | 5472 |
| *Ligands (no.)* | 218 |
| *Solvent (no.)* | 567 |
| *Protein residues (no.)* | 672 |
| *MolProbity score* | 1.11 |
| *Clash score* | 2.79 |
| *Rotamer outliers (%)* | 0.51 |
| *RMS deviation of bond lengths (Å)* | 0.01 |
| *RMS deviation of bond angles (º)* | 1.13 |
| *Ramachandran plot outliers (%)* | 0.00 |
| *Ramachandran plot allowed (%)* | 1.82 |
| *Ramachandran plot favored (%)* | 98.18 |
| *Average B-factors* | 30.07 |
| *macromolecules* | 29.25 |
| *ligands* | 26.38 |
| *solvent* | 38.77 |

The 2mFo-DFc map clearly supports most of the Probe 1 molecule, outlining the spermidine moiety and extending toward the less well-defined propargyl group. Thus, the residual positive mFo-DFc density around the propargyl group may reflect the conformational flexibility of this part of the molecule, which the DHS binding pocket does not strongly coordinate (Fig.3A). Another part of electron density visible deep within the binding site is also present in the crystallographic map of DHS in complex with SPD and NAD (PDB 6XXJ). The authors of that study decided to explain this density by building formic acid (a component of the crystallization buffer) at this position, but the negative mFo-DFc signal is present around this formate in PDB 6XXJ (Fig.3B)(10). Similarly, we observed pronounced negative mFo-DFc density after attempting to fit this map feature with formate. Thus, we placed water at this site. Such an approach generates residual positive mFo-DFc signal in the vicinity (Fig.3A). Hence, this water molecule can likely shift slightly perpendicular to the long axis of Probe 1. The OMIT mFo-DFc map of the ligand and water bound deep within the pocket supports this placement of the derivative (Fig. 3C).

**Figure 3.**
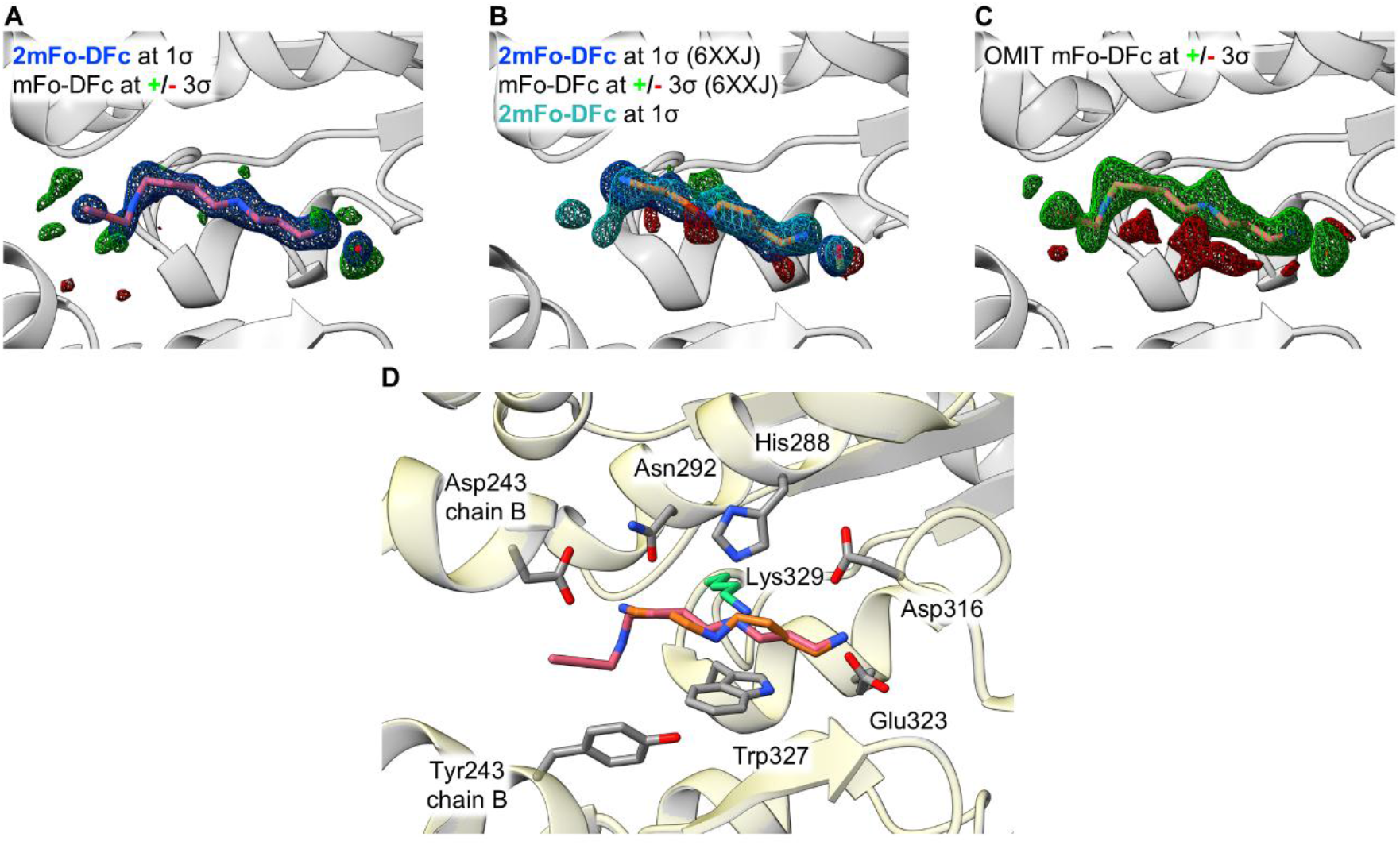
Crystallographic structure of DHS in complex with N8-propargyl spermidine and NAD. **A.** 2mFo-DFc density (blue) of Probe1 (pink) and water bound near N1 of the ligand, buried deep within the binding site, contoured at 1σ. Residual mFo-DFc (positive in green and negative in red) density around the ligand. **B.** 2mFo-DFc density (blue) of spermidine (orange) and formate (yellow) bound deep within the pocket from PDB 6XXJ, superimposed with 2mFo-DFc map (cyan) of Probe 1, contoured at 1σ. Residual mFo-DFc (positive in green and negative in red) density around the ligands from PDB 6XXJ. **C.** OMIT mFo-DFc map (positive in green and negative in red) of Probe 1 (pink) and water bound near N1 of this ligand. **D.** Structural details of Probe 1 (pink) interaction with DHS. The catalytic lysine is shown in green. Superimposed with PDB 6XXJ (light yellow), and SPD shown as orange sticks.

Probe 1 binds to DHS with the N8-propargyl moiety pointing towards the outside of the DHS polyamine binding pocket, in an orientation opposite to that of SPD (Fig.3D). The primary amino group, bound deep within the binding site (N8 of SPD/N1 of Probe 1 - both occupying the exact same position), interacts with Asp316 and Glu323. The middle secondary amine (N4 of SPD) lies between the ε-amine of the catalytic lysine in DHS (Lys329), His288, and Trp327, while N4 of Probe 1, shifted deeper into the pocket, sits slightly closer to the ε-amine of Lys329. N1 of SPD is coordinated by Asn292 as well as Asp243 from the second DHS protomer within the dimer. N8 of Probe 1, localized around N1 of SPD, is located farther from these residues, and the propargyl moiety extending toward the outside of the binding site may interact with Tyr243 from the second DHS protomer within the dimer, which is not an important contributor to SPD binding (Fig.3D). Probe 1 binding did not introduce any conformational changes into DHS when compared with PDB 6XXJ, as the superposition of the molecular models yielded an RMSD for 672 Cα atoms of 0.13 Å (Fig.4D). Overall, these structural data show a non-productive binding pose of Probe 1 incompatible with catalysis and imply that it cannot function as a DHS substrate.

## Discussion

Given our initial interest in using N8-propargyl spermidine (Probe 1) to track DHS-mediated PTMs, we experimentally evaluated its ability to serve as a substrate in the DHS-catalyzed reaction. This question gained particular relevance following a recent study that interpreted proteome-wide labeling with Probe 1 as evidence of widespread protein hypusination and claimed functional consequences of a DHS-dependent modification of RIPK1(29). If independently confirmed, this proposed expansion of DHS substrate specificity would challenge the established view that hypusination is restricted to a single lysine residue in eIF5A.

Using orthogonal approaches, we demonstrated that the widespread labeling generated by N8-propargyl spermidine predominantly reflects CuAAC-associated thiol chemistry rather than DHS-dependent enzymatic transfer. This signal can be largely abolished by cysteine alkylation before click coupling and was independent of DHS activity under all tested conditions. Consistent with these findings, we detected no DHS/DOHH-dependent modification of the canonical substrate eIF5A *in vitro* in the presence of Probe 1. Our findings challenge the recent interpretation of Probe 1-derived labeling as evidence for expanded DHS substrate specificity and highlight important limitations of CuAAC-based workflows employing propargylated polyamines.

The biological properties of natural metabolite cannot be simply assumed to be retained by their tagged derivatives. TvS approaches therefore require experimental validation that the relevant enzyme can accept the analog and transfer its tagged moiety to the intended target. Identifying probe-modified peptides with mass shifts consistent with the expected transfer product would provide particularly important evidence for this reaction. However, we did not detect such peptides in our experiments.

For hypusination, this requirement is especially relevant given the stringent substrate recognition of DHS. Early competitive inhibition studies suggested that compounds capable of binding to DHS must possess at least two charged primary amino or guanidino groups at an appropriate distance(36). In these assays, N8-ethyl spermidine and spermine did not appear to compete with radiolabeled spermidine(36). Nevertheless, later structural studies showed that spermine can occupy the DHS active site, with its additional 3-aminopropyl moiety extending out of the pocket(10), illustrating that binding alone does not establish productive substrate utilization.

Our crystal structure similarly revealed that the N8-propargyl moiety of Probe 1 points out of the polyamine-binding pocket, with the compound adopting an orientation opposite to that of spermidine. This non-productive binding mode provides a structural explanation for the absence of detectable transfer of propargylated 4-aminobutyl group in our experiments. Moreover, it raises the question whether DHS can accommodate N8-acetyl spermidine as a substrate, if direct transfer of an N-acetylated 4-aminobutyl moiety to RIPK1 is considered.

RIPK1 hypusination and K140 acetylhypusination were also reported on the basis of shotgun proteomics and immunoblot analyses beyond the probe-based experiments(29). These assignments would benefit from comparison with synthetic modified peptide standards and further characterization of antibody specificity. The differing modification sites identified *in vitro* and in cells, together with the limited dependence of the reported *in vitro* MS signals on the presence of spermidine, also merit clarification. Evidence from *Dhps*-knockout cells should additionally be examined in light of the broader physiological consequences of impaired eIF5A hypusination(38–41). Alongside these considerations, our inability to detect RIPK1 modification with native spermidine using a well-validated antibody supports the need for further independent verification.

The implications of our results extend beyond Probe 1-derived background chemistry itself. Bioorthogonally tagged analogs of endogenous small molecules, designed for CuAAC-based detection or enrichment, are increasingly used to investigate a broad range of protein post-translational modifications in intact-cell systems, with examples including glycosylation, lipidation, AMPylation, lactylation, and crotonylation(20, 21, 42–46). In such applications, widespread probe-derived labeling cannot be interpreted as evidence of biologically meaningful covalent modification unless rigorously verified. Importantly, in many of these studies, direct identification of probe-modified peptides and modification sites by mass spectrometry constitutes a key line of evidence supporting enzymatic incorporation of the reporter and can have the additional benefit of guiding subsequent site-directed mutagenesis and functional studies.

Our peptide-level analysis of cells metabolically labeled with Probe 1, originally intended to identify sites of enzymatic probe transfer, instead revealed that CuAAC-associated thiol chemistry, consistent with recent reports(27, 35), can generate extensive background labeling that mimics enzymatic probe incorporation. If such side reactions are not adequately controlled, protein-level enrichment alone may therefore lead to erroneous assignment of enzyme substrates, particularly when the existence and chemical identity of the proposed modified amino acid residues are not independently confirmed using rigorously validated orthogonal methods.

Importantly, these observations should not be interpreted as a general limitation of bioorthogonally tagged polyamine analogs as chemical probes. Our recent studies show that polyamine-derived probes can yield selective, biologically informative results when rigorous functional or other validation supports the observed signal (26, 47). For example, polyamine photoaffinity probes enabled mapping of non-covalent polyamine-protein interactions in live cells, including localization of interaction regions, often enriched in acidic amino acids, on several proteins(26). Conversely, such compounds can provide valuable biological insight when their target engagement is carefully substantiated.

Taken together, our findings cast substantial doubt on the existence of an extensive hypusinated proteome. The widespread labeling observed with Probe 1 can be largely explained by CuAAC-associated thiol chemistry rather than enzymatic modification, undermining the interpretation of large-scale protein enrichment as evidence for numerous DHS substrates. Our results highlight the importance of experimental confirmation with complementary methods when interpreting CuAAC-based proteomic analyses before proposing expanded substrate specificity for PTM-generating enzymes.

### Experimental procedures

#### Synthesis of N8-propargyl spermidine (Probe 1)

Probe 1 was synthesized according to the procedure previously reported(28). Detailed synthetic methods and analytical characterization data of the preparation used in this study, including NMR and HRMS spectra, are available in the same publication(28).

#### Cell lines and culture conditions

HEK293T and MEF cells were cultured in high-glucose DMEM supplemented with fetal bovine serum (10%), penicillin (100 U/mL), and streptomycin (100 µg/mL) at 37 °C in a humidified atmosphere of 5% CO₂ and 95% air. Cell lines were regularly tested for mycoplasma contamination using a PCR-based method.

#### Treatment in cells

HEK293T or MEF cells were treated with Probe 1 or SPD (200-400 µM) for 1-4 h. Where indicated, cells were pretreated with GC-7 (40-56 μM) for 1-4 h. After washing, cells were lysed in 2% SDS/PBS. Where indicated, lysates were treated with iodoacetamide (40 mM; IAA) for 30 min. Protein concentrations were determined by DC Protein Assay and adjusted to 1 mg/mL in 0.5% SDS/PBS. Protein samples were then supplemented with 5-TAMRA-azide (0.1 mM) or azido-PEG₃-biotin (0.2 mM), along with THPTA (0.2 mM), CuSO₄ (1 mM), and TCEP (1 mM). Samples were vortex-mixed for 1 h at RT, and CuAAC was quenched with EDTA (5 mM). Methanol (1 vol.) and chloroform (0.25 vol.) were added, and the samples were briefly vortex-mixed and centrifuged at 21,000 g for 5 min to pellet precipitated proteins. The pellets were washed with methanol and dried. Detailed descriptions of the experimental procedures are available in the Supporting Information.

#### Sample processing for in-gel fluorescence after treatment in cells

After ligation with 5-TAMRA-azide and protein precipitation, the pellets were boiled in sample loading buffer containing β-mercaptoethanol (10%). Proteins were separated by SDS-PAGE (12% polyacrylamide gels). In-gel fluorescence was recorded using an Amersham Imager, and total protein was subsequently visualized by Coomassie Brilliant Blue staining.

#### Sample processing for proteomics after treatment in cells (protein-level enrichment of biotinylated proteins)

After ligation with azido-PEG3-biotin and protein precipitation from MEF lysates, the pellets were resuspended in 2% SDS/PBS, diluted to 0.5% SDS, and incubated with NeutrAvidin agarose for 2 h at RT. The beads were washed and resuspended in HEPES (100 mM, pH 8.0) containing TCEP (5 mM) and chloroacetamide (10 mM; CAA). Sequencing Grade Modified Trypsin was added, and the samples were incubated overnight at 37 °C. The supernatants were collected and acidified with trifluoroacetic acid (1%), and the resulting tryptic peptides were labeled using an on-column TMT labeling protocol(48). LC–MS/MS analysis was performed using an UltiMate 3000 nano-LC system coupled to a Q Exactive HF-X mass spectrometer via an Easy-Spray source. Peptides were separated on an Easy-Spray Acclaim PepMap C18 column using an acetonitrile gradient. LC– MS/MS data were processed in MaxQuant using the reviewed mouse proteome from UniProt and standard settings(49). In Perseus(50), standard filters were applied, and corrected reporter ion intensities were log₂-transformed. Proteins enriched in the Probe 1 condition relative to the spermidine condition were identified using a one-sided Student’s t-test (permutation-based FDR = 0.01, S₀ = 0.5, n = 3). The results were exported from Perseus and formatted in Microsoft Excel. Detailed descriptions of the experimental procedures are available in the Supporting Information.

#### Sample processing for proteomics after treatment in cells (peptide-level enrichment of biotinylated peptides)

After ligation with azido-PEG₃-biotin and protein precipitation from HEK293T lysates, the pellets were resuspended in HEPES (100 mM, pH 8.0) containing TCEP (5 mM) and CAA (10 mM). Samples were digested overnight at 37 °C with Sequencing Grade Modified Trypsin. Residual trypsin activity was quenched by adding EDTA-free Protease Inhibitor Cocktail, samples were cleared by centrifugation, and incubated with NeutrAvidin agarose for 2 h at RT. The beads were washed, and probe-modified peptides eluted with 0.1% trifluoroacetic acid and 0.2% formic acid in 80% acetonitrile at 37 °C. After pre-cleaning on C18 StageTips, the peptides were subjected to LC-MS/MS analysis. LC–MS/MS data were processed in MaxQuant using the UniProt human reference proteome and standard settings, with three custom variable modifications included in the search. These were the triazole-tagged forms of Probe 1-derived hypusine (Δmass = 569.2996 Da) and deoxyhypusine (Δmass = 553.3046 Da), and the cysteine–Probe 1 thiotriazole adduct (Δmass = 625.3734 Da). No peptides carrying the biotinylated triazole-tagged forms of Probe 1-derived hypusine or deoxyhypusine were identified. The results for peptides carrying the cysteine–Probe 1 thiotriazole adduct were filtered in Perseus using standard filters, then exported and formatted in Microsoft Excel. Detailed descriptions of the experimental procedures are available in the Supporting Information.

#### Native cell lysate-based in-gel fluorescence analysis

Confluent culture of HEK293T cells from a T75 flask was washed with PBS, and lysed in 1ml of buffer (20 mM Tris pH 8.0, 100 mM NaCl, 0.5% NP-40, Pierce EDTA-free protease inhibitor cocktail). The lysate was clarified by centrifugation (15,000 x g, 20 min, 4°C). Protein concentration in the supernatant was measured using a NanoDrop 2000, assuming A_280_=1 corresponds to 1 mg/ml protein. The (deoxy)hypusination reaction was carried out by incubating 120 µg of the lysate with or without 1 mM of GC-7 (mixed with the lysate 15 min before any other reagents were added to the reaction mixture), 12 µg of recombinant human DHS (in samples containing recombinant DHS, 1 mM NAD was also included), and 0.5 mM Probe 1 in PBS (total reaction volume 100 µl) for 1 h at RT. Samples were then reacted or not with 40 mM IAA for 30 min at RT. Proteins were precipitated by the addition of methanol (1 vol.) and chloroform (1/4 vol.), as described earlier. Protein pellets were washed with methanol, air-dried, and reconstituted in 25 µl 2% SDS in PBS. Then 75 µl of PBS was added, and the samples underwent CuAAC ligation with azide-functionalized TAMRA, as described earlier under treatment in cells. The samples were processed as described in the sample processing for in-gel fluorescence after treatment in cells, but the in-gel fluorescence was recorded using a ChemiDoc MP in the Cy3 channel.

#### Recombinant protein purification

Plasmids used for the production of recombinant human DHS and eIF5A were described before(12). The DNA encoding human DOHH (UniProt: Q9BU89, residues 1–289) was synthesized along with an N-terminal TEV site and cloned into the pGEX-6P-1 vector (BamHI/XhoI; Genscript). The DNA encoding the kinase domain of human RIPK1 (UniProt: Q13546, residues 1–307) was synthesized along with an N-terminal 6xHis-tag followed by a TEV site and cloned into the pET-24d(+) vector (NcoI/SacI; Genscript). All proteins were expressed and purified according to a protocol described previously(12). For DOHH, the clarified lysate was loaded onto a GSTrap FF 5 ml column (Cytiva), and the column was washed with 20 CV of buffer (50 mM Tris-HCl pH 7.8, 200 mM NaCl, 5% glycerol, 3 mM β-ME). The protein was eluted with the same buffer containing 30 mM reduced glutathione. All subsequent steps were as described previously, but we isolated the tag-free protein using reverse GSTrap FF column chromatography. Proper folding of the recombinant RIPK1 was verified by analysis of its autophosphorylation(51) using the ATPγS-based assay described previously(12). Briefly, 2 ug of the protein was incubated in 30 µl of buffer (20 mM HEPES pH 7.3, 10 mM MnCl2, 10 mM MgCl2, 150 mM NaCl) supplemented with 1 mM of ATPyS for 1 h at RT, and after reaction with 50 mM of p-nitrobenzylmesylate samples were mixed with SDS-PAGE sample buffer and used for Western Blot analysis with detection using anti-thiophosphate ester antibody (Abcam, ab92570) (Fig.S1).

#### LC-MS/MS analysis of human recombinant eIF5A after *in vitro* reaction with DHS/DOHH

eIF5A (20 µg) was incubated for 1 h at RT in glycine/NaOH buffer (0.2 M) containing NAD (1 mM) and either SPD (1 mM) or Probe 1 (1 mM), with or without DHS (2 µg) and DOHH (2 µg). Samples were diluted with HEPES buffer (100 mM, pH 8.0) and digested with trypsin overnight at 37 °C in the presence of TCEP (5 mM) and CAA (10 mM). Peptides were cleaned up using C18 StageTips before LC–MS/MS analysis. LC–MS/MS data were processed in MaxQuant using standard settings with four custom variable modifications and the UniProt *E. coli* reference proteome supplemented with the sequences of human DOHH, DHS, and eIF5A. The mass shifts were 87.0684 Da for hypusine, 71.0735 Da for deoxyhypusine, and 125.0841 Da / 109.0892 Da for their respective propargylated forms. The MaxQuant modificationSpecificPeptides output was imported into Perseus, where eIF5A peptides were retained, and their intensities were log₂-transformed. Unmodified peptides that did not span Lys50 were retained only if intensities were available across all compared conditions. For unmodified peptides spanning Lys50, missing intensities in complete reactions containing both DHS and DOHH were imputed from a normal distribution. Within each sample, unmodified peptide intensities were median-centered on the log₂ scale. Modified peptide intensities were analysed without imputation or median centering. The results were exported to Microsoft Excel for final formatting. Detailed descriptions of the experimental procedures are available in the Supporting Information.

#### Western blot-based *in vitro* (deoxy)hypusination assay

The *in vitro* reaction was carried out as described(29). 2 μg RIPK1 or eIF5A was incubated with or without 0.2 μg DHS, 0.2 μg DOHH, and 1 mM spermidine in 40 μl of 0.2 M glycine-NaOH pH 9.2 buffer with 1 mM DTT and 1 mM NAD. The reaction was stopped by the addition of 8 μl 6x SDS-PAGE sample buffer and boiled for 5 min. 20 μL of each sample was separated on 12% SDS-PAGE gel for immunoblotting. Western Blot detection with context-independent anti-(deoxy)hypusine antibody (Creative Biolabs, PABL-582) was performed as previously described (12).

#### Protein crystallization

DHS was co-crystallized with Probe 1, as described previously(10). Briefly, equal amounts of protein solution (∼18 mg/ml) supplemented with 1 mM NAD and 10 mM Probe 1, and crystallization buffer (170 mM carboxylic acid mix and 47.5% precipitant mix (parts of the Morpheus Screen from the MRC Laboratory of Molecular Biology), 100 mM Tris-Bicine, pH 8.5) were mixed and left to equilibrate by the sitting drop vapor diffusion technique at RT. Crystals were harvested using cryo-loops and flash-frozen in liquid nitrogen.

#### Diffraction data collection and structure determination

X-ray diffraction data were collected under cryogenic conditions at the BM07 - FIP2 beamline operated by the ESRF. The collected data were processed using XDSAPP3(52). We solved the structure by molecular replacement using PDB 6XXJ (with all ligands except NAD removed) as the search model, followed by initial refinement in DIMPLE (53). The model was manually rebuilt in *Coot*(54). Ligand restraints were generated with eLBOW(55). The structure was refined using Phenix Refine(56). Water molecules were automatically placed during refinement and subsequently manually inspected in Coot. The OMIT map was generated with the Phenix suite. The final model was validated with MolProbity(57). All visualizations of molecular models and maps were prepared using UCSF ChimeraX(58).

## Data availability

LC-MS/MS data have been deposited to the ProteomeXchange Consortium via the PRIDE(59) partner repository with the dataset identifiers PXD080059 and PXD071577. The crystallographic structure has been deposited in the Protein Data Bank with the accession code 35PO.

## Supporting information

This article contains supporting information.

## Acknowledgements

Proteomic measurements were performed at the Proteomics Core Facility, IMol Polish Academy of Sciences. We extend our gratitude to Mrs. Viktoriia Lastivka for LC-MS/MS measurements. We acknowledge the MCB Structural Biology Core Facility for access to instruments and especially Ms. Klaudia Woś for technical support with protein crystallization, and the European Synchrotron Radiation Facility (ESRF) for provision of synchrotron radiation facilities under proposal MX 2791 on beamline BM07 - FIP2 (supported by the French ANR PIA3 (France 2030) EquipEx+ project MAGNIFIX under grant agreement ANR-21-ESRE-0011) and the staff of the ESRF for assistance and support during the beamtime. We thank Prof. Agnieszka Chacińska (IMol) and Dr. Karolina Szczepanowska (IMol) for providing the HEK293T and MEF cells, respectively. Text editing was supported using the OpenAI ChatGPT-6 environment.

## Author contributions

Conceptualization, P.K., P.G. and R.S.; methodology, P.K. and R.S; MS data analysis and curation, R.S.; chemical synthesis P.W.; *in cellulo* experiments, M.C.; *in vitro* experiments, P.K.; protein crystallography, M.S.-S., P.K.; figure preparation, P.K., M.C. and R.S.; writing – original draft preparation, P.K. and R.S.; writing – review & editing, P.K., P.G. and R.S; supervision, P.G. and R.S.

## Funding and additional information

This research was funded by the National Science Centre, Poland grants: UMO-2020/38/E/ST4/00250 (to R.S.), UMO-2024/54/E/NZ1/00265 (to P.G.) and UMO-2023/49/N/NZ1/03559 (to P.K.). P.K. was supported by the Foundation for Polish Science (FNP) - START scholarship. Access to the ESRF was financed by the Polish Ministry of Science and Higher Education – decision no. 2021/WK/11.

## Conflict of interest

The authors declare that they have no conflicts of interest with the contents of this article.

## Notes

https://www.ebi.ac.uk/pride/archive/projects/PXD071577

https://www.ebi.ac.uk/pride/archive/projects/PXD080059

### Competing Interest Statement

The authors have declared no competing interest.

## References

1. Joe, Y. A., Wolff, E. C., and Park, M. H. (1995) Cloning and Expression of Human Deoxyhypusine Synthase cDNA. Journal of Biological Chemistry. 270, 22386–22392

2. Park, M. H., Cooper, H. L., and Folk, J. E. (1982) The biosynthesis of protein-bound hypusine (N epsilon -(4-amino-2-hydroxybutyl)lysine). Lysine as the amino acid precursor and the intermediate role of deoxyhypusine (N epsilon -(4-aminobutyl)lysine). J Biol Chem. 257, 7217–7222

3. Park, M. H., Liberato, D. J., Yergey, A. L., and Folk, J. E. (1984) The biosynthesis of hypusine (N epsilon-(4-amino-2-hydroxybutyl)lysine). Alignment of the butylamine segment and source of the secondary amino nitrogen. J Biol Chem. 259, 12123–12127

4. Park, M. H., and Wolff, E. C. (1988) Cell-free synthesis of deoxyhypusine. Separation of protein substrate and enzyme and identification of 1,3-diaminopropane as a product of spermidine cleavage. Journal of Biological Chemistry. 263, 15264–15269

5. Han, Z., Sakai, N., Böttger, L. H., Klinke, S., Hauber, J., Trautwein, A. X., and Hilgenfeld, R. (2015) Crystal Structure of the Peroxo-diiron(III) Intermediate of Deoxyhypusine Hydroxylase, an Oxygenase Involved in Hypusination. Structure. 23, 882–892

6. Zakaszewski, D., Wilk, P., Wątor-Wilk, E., and Grudnik, P. (2026) Mismetallation reveals metal tolerance and enhanced stability of human deoxyhypusine hydroxylase. Int J Biol Macromol. 384, 154732

7. Park, M. H., Cooper, H. L., and Folk, J. E. (1981) Identification of hypusine, an unusual amino acid, in a protein from human lymphocytes and of spermidine as its biosynthetic precursor. Proc. Natl. Acad. Sci. U.S.A. 78, 2869–2873

8. Cooper, H. L., Park, M. H., Folk, J. E., Safer, B., and Braverman, R. (1983) Identification of the hypusine-containing protein hy+ as translation initiation factor eIF-4D. Proc. Natl. Acad. Sci. U.S.A. 80, 1854–1857

9. Wolff, E. C., Kang, K. R., Kim, Y. S., and Park, M. H. (2007) Posttranslational synthesis of hypusine: evolutionary progression and specificity of the hypusine modification. Amino Acids. 33, 341–350

10. Wątor, E., Wilk, P., and Grudnik, P. (2020) Half Way to Hypusine-Structural Basis for Substrate Recognition by Human Deoxyhypusine Synthase. Biomolecules. 10, 522

11. Wątor-Wilk, E., Wilk, P., and Grudnik, P. (2025) The structural biology of deoxyhypusination complexes. Structure. 33, 221–227

12. Becker, A. E., Kochanowski, P., Wu, P.-K., Wątor, E., Chen, W., Guchhait, K., Biela, A. P., Grudnik, P., and Park, J.-I. (2024) ERK1/2 interaction with DHPS regulates eIF5A deoxyhypusination independently of ERK kinase activity. Cell Rep. 43, 114831

13. Wątor, E., Wilk, P., Biela, A., Rawski, M., Zak, K. M., Steinchen, W., Bange, G., Glatt, S., and Grudnik, P. (2023) Cryo-EM structure of human eIF5A-DHS complex reveals the molecular basis of hypusination-associated neurodegenerative disorders. Nat Commun. 14, 1698

14. Cooper, H. L., Park, M. H., and Folk, J. E. (1982) Posttranslational formation of hypusine in a single major protein occurs generally in growing cells and is associated with activation of lymphocyte growth. Cell. 29, 791– 797

15. Joe, Y. A., and Park, M. H. (1994) Structural features of the eIF-5A precursor required for posttranslational synthesis of deoxyhypusine. J Biol Chem. 269, 25916–25921

16. Thompson, G. M., Cano, V. S. P., and Valentini, S. R. (2003) Mapping eIF5A binding sites for Dys1 and Lia1: in vivo evidence for regulation of eIF5A hypusination. FEBS Lett. 555, 464–468

17. Kho, Y., Kim, S. C., Jiang, C., Barma, D., Kwon, S. W., Cheng, J., Jaunbergs, J., Weinbaum, C., Tamanoi, F., Falck, J., and Zhao, Y. (2004) A tagging-via-substrate technology for detection and proteomics of farnesylated proteins. Proc Natl Acad Sci U S A. 101, 12479–12484

18. Rostovtsev, V. V., Green, L. G., Fokin, V. V., and Sharpless, K. B. (2002) A stepwise huisgen cycloaddition process: copper(I)-catalyzed regioselective “ligation” of azides and terminal alkynes. Angew Chem Int Ed Engl. 41, 2596–2599

19. Tornøe, C. W., Christensen, C., and Meldal, M. (2002) Peptidotriazoles on Solid Phase: [1,2,3]-Triazoles by Regiospecific Copper(I)-Catalyzed 1,3-Dipolar Cycloadditions of Terminal Alkynes to Azides. J. Org. Chem. 67, 3057–3064

20. Storck, E. M., Morales-Sanfrutos, J., Serwa, R. A., Panyain, N., Lanyon-Hogg, T., Tolmachova, T., Ventimiglia, L. N., Martin-Serrano, J., Seabra, M. C., Wojciak-Stothard, B., and Tate, E. W. (2019) Dual chemical probes enable quantitative system-wide analysis of protein prenylation and prenylation dynamics. Nat Chem. 11, 552– 561

21. Thinon, E., Serwa, R. A., Broncel, M., Brannigan, J. A., Brassat, U., Wright, M. H., Heal, W. P., Wilkinson, A. J., Mann, D. J., and Tate, E. W. (2014) Global profiling of co- and post-translationally N-myristoylated proteomes in human cells. Nat Commun. 5, 4919

22. Serwa, R. A., Abaitua, F., Krause, E., Tate, E. W., and O’Hare, P. (2015) Systems Analysis of Protein Fatty Acylation in Herpes Simplex Virus-Infected Cells Using Chemical Proteomics. Chem Biol. 22, 1008–1017

23. Wilson, J. P., Raghavan, A. S., Yang, Y.-Y., Charron, G., and Hang, H. C. (2011) Proteomic analysis of fatty-acylated proteins in mammalian cells with chemical reporters reveals S-acylation of histone H3 variants. Mol Cell Proteomics. 10, M110.001198

24. Martin, B. R., Wang, C., Adibekian, A., Tully, S. E., and Cravatt, B. F. (2011) Global profiling of dynamic protein palmitoylation. Nat Methods. 9, 84–89

25. DeGraw, A. J., Palsuledesai, C., Ochocki, J. D., Dozier, J. K., Lenevich, S., Rashidian, M., and Distefano, M. D. (2010) Evaluation of alkyne-modified isoprenoids as chemical reporters of protein prenylation. Chem Biol Drug Des. 76, 460–471

26. Zakrzewski, M., Sas, Z., Cocom-Chan, B., Firdaus, M. E. R., Kałek, M., Szczepanowska, K., Gerlach, P., Marusiak, A., and Serwa, R. A. (2025) Profiling polyamine–protein interactions in live cells through photoaffinity labeling. *RSC Chem*. Biol. 6, 1787–1796

27. Sas, Z., Zakrzewski, M., Wardak, A. D., Wanat, P., Czestkowski, W., Zmorzyńska, J., Marusiak, A. A., and Serwa, R. A. (2026) Minimal-tag polyamines for *in situ* probing of their intracellular distribution. Chem. Commun. 62, 1623–1627

28. Tran, T. T. B., Sas, Z., Karpińska, K., Wanat, P., Marusiak, A. A., and Serwa, R. A. (2026) Spermidine-Based Activity Probe for ALDH1A1 in Cells. Chembiochem. 27, e70399

29. Zhang, T., Fu, W., Zhang, H., Li, J., Xing, B., Cai, Y., Zhang, M., Liu, X., Qi, C., Qian, L., Hu, X., Zhu, H., Yang, S., Zhang, M., Liu, J., Li, G., Li, Y., Xiang, R., Qi, Z., Hu, J., Li, Y., Zou, C., Wang, Q., Jin, X., Pang, R., Li, P., Liu, J., Zhang, Y., Wang, Z., Zhu, Z.-J., Shan, B., and Yuan, J. (2024) Spermidine mediates acetylhypusination of RIPK1 to suppress diabetes onset and progression. Nat Cell Biol. 26, 2099–2114

30. Degterev, A., Hitomi, J., Germscheid, M., Ch’en, I. L., Korkina, O., Teng, X., Abbott, D., Cuny, G. D., Yuan, C., Wagner, G., Hedrick, S. M., Gerber, S. A., Lugovskoy, A., and Yuan, J. (2008) Identification of RIP1 kinase as a specific cellular target of necrostatins. Nat Chem Biol. 4, 313–321

31. Silke, J., Rickard, J. A., and Gerlic, M. (2015) The diverse role of RIP kinases in necroptosis and inflammation. Nat Immunol. 16, 689–697

32. Liu, Z., and Chan, F. K.-M. (2021) Regulatory mechanisms of RIPK1 in cell death and inflammation. Seminars in Cell & Developmental Biology. 109, 70–75

33. Delanghe, T., Dondelinger, Y., and Bertrand, M. J. M. (2020) RIPK1 Kinase-Dependent Death: A Symphony of Phosphorylation Events. Trends in Cell Biology. 30, 189–200

34. Umland, T. C., Wolff, E. C., Park, M. H., and Davies, D. R. (2004) A new crystal structure of deoxyhypusine synthase reveals the configuration of the active enzyme and of an enzyme.NAD.inhibitor ternary complex. J Biol Chem. 279, 28697–28705

35. Wiest, A., and Kielkowski, P. (2024) Cu-Catalyzed Azide-Alkyne-Thiol Reaction Forms Ubiquitous Background in Chemical Proteomic Studies. J Am Chem Soc. 146, 2151–2159

36. Jakus, J., Wolff, E. C., Park, M. H., and Folk, J. E. (1993) Features of the spermidine-binding site of deoxyhypusine synthase as derived from inhibition studies. Effective inhibition by bis- and mono-guanylated diamines and polyamines. Journal of Biological Chemistry. 268, 13151–13159

37. Zhai, Q., He, M., Song, A., Deshayes, K., Dixit, V. M., and Carter, P. J. (2016) Structural Analysis and Optimization of Context-Independent Anti-Hypusine Antibodies. J Mol Biol. 428, 603–617

38. Ganapathi, M., Padgett, L. R., Yamada, K., Devinsky, O., Willaert, R., Person, R., Au, P.-Y. B., Tagoe, J., McDonald, M., Karlowicz, D., Wolf, B., Lee, J., Shen, Y., Okur, V., Deng, L., LeDuc, C. A., Wang, J., Hanner, A., Mirmira, R. G., Park, M. H., Mastracci, T. L., and Chung, W. K. (2019) Recessive Rare Variants in Deoxyhypusine Synthase, an Enzyme Involved in the Synthesis of Hypusine, Are Associated with a Neurodevelopmental Disorder. Am J Hum Genet. 104, 287–298

39. Nishimura, K., Lee, S. B., Park, J. H., and Park, M. H. (2012) Essential role of eIF5A-1 and deoxyhypusine synthase in mouse embryonic development. Amino Acids. 42, 703–710

40. Padgett, L. R., Robertson, M. A., Anderson-Baucum, E. K., Connors, C. T., Wu, W., Mirmira, R. G., and Mastracci, T. L. (2021) Deoxyhypusine synthase, an essential enzyme for hypusine biosynthesis, is required for proper exocrine pancreas development. FASEB J. 35, e21473

41. Sasaki, K., Abid, R., and Miyazaki, M. (1996) Deoxyhypusine synthase gene is essential for cell viability in the yeast *Saccharomyces cerevisiae*. FEBS Letters. 384, 151–154

42. Sun, Y., Chen, Y., and Peng, T. (2022) A bioorthogonal chemical reporter for the detection and identification of protein lactylation. Chem Sci. 13, 6019–6027

43. Zaro, B. W., Yang, Y.-Y., Hang, H. C., and Pratt, M. R. (2011) Chemical reporters for fluorescent detection and identification of O-GlcNAc-modified proteins reveal glycosylation of the ubiquitin ligase NEDD4-1. Proc Natl Acad Sci U S A. 108, 8146–8151

44. Martin, B. R., and Cravatt, B. F. (2009) Large-scale profiling of protein palmitoylation in mammalian cells. Nat Methods. 6, 135–138

45. Zhou, Y.-F., Yuan, S., Ma, B., Gao, J., and Wang, C. (2025) Chemical proteomic profiling of lysine crotonylation using minimalist bioorthogonal probes in mammalian cells. Chem Sci. 16, 2843–2849

46. Kielkowski, P., Buchsbaum, I. Y., Kirsch, V. C., Bach, N. C., Drukker, M., Cappello, S., and Sieber, S. A. (2020) FICD activity and AMPylation remodelling modulate human neurogenesis. Nat Commun. 11, 517

47. Zabala-Letona, A., Pujana-Vaquerizo, M., Martinez-Laosa, B., Ponce-Rodriguez, M., Garcia-Longarte, S., Mendizabal, I., Gimeno, A., Rogalska, M., Tan, J., Cabrera, D., van Liempd, S., Ximenez-Embun, P., Espinosa, S., Fagoaga-Eugui, M., Peccati, F., Zakrzewski, M., Astobiza, I., Arana-Castañares, M., Cherkaoui, S., Sendino, M., Martín-Barros, I., Ercilla, A., Bozal-Basterra, L., Carlevaris, O., Arruabarrena-Aristorena, A., Pérez-Andrés, E., Santamaría-Zamorano, T., Ferrer-Bonsoms, J. A., Carazo, F., Cieśla, M., Lobato, C., Seoane, J., Martín-Martín, N., Barrio, R., Sutherland, J. D., Aransay, A. M., Falcón-Pérez, J. M., Martínez-Pastor, B., Rubio, A., Blanco, F. J., Hogarty, M. D., Morscher, R. J., Berra, E., Serwa, R. A., Jiménez-Barbero, J., Jiménez-Osés, G., Efeyan, A., Finley, L., Lizcano, J. M., Muñoz, J., Valcarcel, J., and Carracedo, A. (2026) Polyamine-dependent metabolic shielding regulates alternative splicing. Nature. 651, 819–828

48. Myers, S. A., Rhoads, A., Cocco, A. R., Peckner, R., Haber, A. L., Schweitzer, L. D., Krug, K., Mani, D. R., Clauser, K. R., Rozenblatt-Rosen, O., Hacohen, N., Regev, A., and Carr, S. A. (2019) Streamlined Protocol for Deep Proteomic Profiling of FAC-sorted Cells and Its Application to Freshly Isolated Murine Immune Cells. Mol Cell Proteomics. 18, 995–1009

49. Tyanova, S., Temu, T., and Cox, J. (2016) The MaxQuant computational platform for mass spectrometry-based shotgun proteomics. Nat Protoc. 11, 2301–2319

50. Tyanova, S., Temu, T., Sinitcyn, P., Carlson, A., Hein, M. Y., Geiger, T., Mann, M., and Cox, J. (2016) The Perseus computational platform for comprehensive analysis of (prote)omics data. Nat Methods. 13, 731–740

51. McQuade, T., Cho, Y., and Chan, F. K.-M. (2013) Positive and negative phosphorylation regulates RIP1- and RIP3-induced programmed necrosis. Biochem J. 456, 409–415

52. Krug, M., Weiss, M. S., Heinemann, U., and Mueller, U. (2012) *XDSAPP* : a graphical user interface for the convenient processing of diffraction data using *XDS*. J Appl Crystallogr. 45, 568–572

53. Wojdyr, M., Keegan, R., Winter, G., and Ashton, A. (2013) DIMPLE - a pipeline for the rapid generation of difference maps from protein crystals with putatively bound ligands

54. Emsley, P., and Cowtan, K. (2004) *Coot* : model-building tools for molecular graphics. Acta Crystallogr D Biol Crystallogr. 60, 2126–2132

55. Moriarty, N. W., Grosse-Kunstleve, R. W., and Adams, P. D. (2009) electronic Ligand Builder and Optimization Workbench (eLBOW): a tool for ligand coordinate and restraint generation. Acta Crystallogr D Biol Crystallogr. 65, 1074–1080

56. Adams, P. D., Afonine, P. V., Bunkóczi, G., Chen, V. B., Davis, I. W., Echols, N., Headd, J. J., Hung, L.-W., Kapral, G. J., Grosse-Kunstleve, R. W., McCoy, A. J., Moriarty, N. W., Oeffner, R., Read, R. J., Richardson, D. C., Richardson, J. S., Terwilliger, T. C., and Zwart, P. H. (2010) *PHENIX* : a comprehensive Python-based system for macromolecular structure solution. Acta Crystallogr D Biol Crystallogr. 66, 213–221

57. Williams, C. J., Headd, J. J., Moriarty, N. W., Prisant, M. G., Videau, L. L., Deis, L. N., Verma, V., Keedy, D. A., Hintze, B. J., Chen, V. B., Jain, S., Lewis, S. M., Arendall, W. B., Snoeyink, J., Adams, P. D., Lovell, S. C., Richardson, J. S., and Richardson, D. C. (2018) MolProbity: More and better reference data for improved all-atom structure validation. Protein Science. 27, 293–315

58. Pettersen, E. F., Goddard, T. D., Huang, C. C., Meng, E. C., Couch, G. S., Croll, T. I., Morris, J. H., and Ferrin, T. E. (2021) UCSF CHIMERAX : Structure visualization for researchers, educators, and developers. Protein Science. 30, 70–82

59. Perez-Riverol, Y., Bandla, C., Kundu, D. J., Kamatchinathan, S., Bai, J., Hewapathirana, S., John, N. S., Prakash, A., Walzer, M., Wang, S., and Vizcaíno, J. A. (2025) The PRIDE database at 20 years: 2025 update. Nucleic Acids Res. 53, D543–D553

