## Supporting Information for "Evaluation of N8-propargyl spermidine as a chemical probe for deoxyhypusine synthase-dependent post-translational modifications"

1. **Table S1.** Results of LC-MS/MS analysis comparing protein level enrichment from MEF cells treated with Probe 1 or SPD after CuAAC ligation to biotin
2. **Table S2.** Results of LC-MS/MS analysis of Probe 1-modified peptides after CuAAC ligation to biotin from HEK293T cells, identifying cysteine-linked thiotriazole adducts
3. **Table S3.** Results of LC-MS/MS analyses of eIF5A peptides after *in vitro* (deoxy)hypusination reaction with Probe 1 or SPD
4. **Figure S1.** Purification of recombinant RIPK1 kinase domain

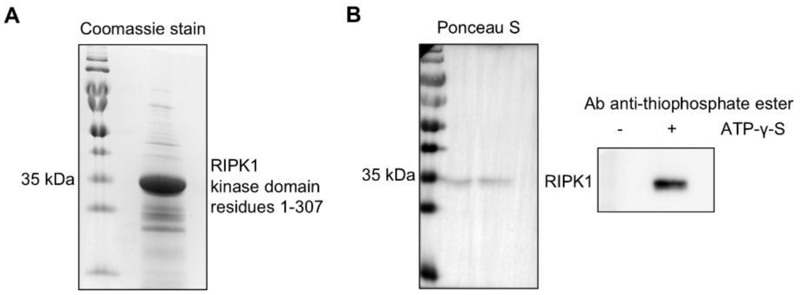


**Figure S1. Purification of recombinant RIPK1 kinase domain A.** Purity of the recombinant RIPK1 kinase domain assessed by SDS-PAGE followed by Coomassie Brilliant Blue staining. **B.** Functional validation of the recombinant RIPK1 kinase domain by assessing its autophosphorylation using an ATPγS-based assay.

**Detailed descriptions of the experimental procedures**

Treatment in cells

HEK293T or MEF cells were washed with PBS and incubated with Probe 1 or spermidine (200-400 µM) for 1-4 h. Where indicated, cells were pretreated with GC7 (40-56 µM) for 1-4 h. Cells were washed with PBS twice, lysed in 2% SDS in PBS, and probe sonicated (3 pulses of 5 s). Lysates were treated where indicated with iodoacetamide (40 mM) for 30 min to prevent the azide–alkyne–thiol reaction, and proteins were then precipitated by adding methanol (1 vol.) and chloroform (0.25 vol.) and briefly vortexing the samples. The pellets were washed with methanol twice, air-dried, and reconstituted in 2% SDS in PBS. The solutions were diluted with PBS to the final concentration of 0.5% SDS and the protein concentration was determined by DC Protein Assay (Bio-Rad). Protein concentrations were adjusted to 1 mg/ml using 0.5% SDS in PBS, and the samples were subjected to CuAAC ligation. A click reaction mixture was prepared by combining 4 reagents (volumes given per 100 µg of proteins): 5-TAMRA-azide (Jena Bioscience, 1 µl, stock 10 mM in DMSO, final conc. 0.1 mM) or azido-PEG3-biotin (Thermo Fisher Scientific, 1 µl, stock 20 mM in water, final conc. 0.2 mM), CuSO4 (2 µl, stock 50 mM in water, final conc. 1 mM), tris(2-carboxyethyl)phosphine (TCEP) (2 µl, stock 50 mM in water, final conc. 1 mM), tris((1-hydroxy-propyl-1H-1,2,3-triazol-4-yl)methyl)amine (THPTA) (1 µl, stock 20 mM in water, final conc. 0.2 mM). Click mixture (6 µl) was added to protein samples (100 µg). The samples were vortex-mixed at RT for 1 h. EDTA (final conc. 5 mM), methanol (1 vol.), and chloroform (1/4 vol.) were added to each sample. The samples were quickly vortexed and centrifuged at 21,000 g for 5 min to pellet precipitated proteins. The pellets were washed with methanol twice and dried.

Sample processing for proteomics after treatment in cells (protein-level enrichment of biotinylated proteins)

Following the ligation in solution to azido-PEG$_3$-biotin in MEF lysates and protein precipitation, the pellets were resuspended in 2% SDS in PBS and PBS was added to dilute the sample (final concentration 1 mg ml$^{-1}$, 0.5% SDS). NeutrAvidin agarose resin was washed with 0.5% SDS in PBS (3×). The samples were added to the beads, and the enrichment was carried out for 2 h at RT with 1100 RPM shaking. The supernatants were removed and the beads were washed with 0.5% SDS in PBS (2×), with PBS (2×), and then with 100 mM HEPES pH 8.0 (2×). The beads were suspended in 100 mM HEPES pH 8.0, containing 5 mM TCEP and 10 mM chloroacetamide. Sequencing Grade Modified Trypsin (Promega) was added and the samples were incubated overnight at 37 °C with 1100 RPM shaking. The samples were centrifuged and the supernatants were transferred into clean tubes. The solutions were acidified with trifluoroacetic acid (TFA, to final conc. 1%) and tryptic peptides were labelled using an on-column TMT labelling protocol. TMT-labelled peptides were compiled into a single TMT sample and concentrated using a SpeedVac concentrator. Prior to the LC-MS/MS measurements, the peptide sample was resuspended in 0.1% TFA and 2% acetonitrile in water. The sample was injected twice. Chromatographic separation was performed on an Easy-Spray Acclaim PepMap column (50 cm length × 75 µm inner diameter, Thermo Fisher Scientific) at 55 °C by applying 180 min acetonitrile gradients in 0.1% aqueous formic acid at a flow rate of 300 nl/min. An UltiMate 3000 nano-LC system was coupled to a Q Exactive HF-X mass spectrometer via an easy-spray source (all Thermo Fisher Scientific). The Q Exactive HF-X was operated in TMT mode with survey scans acquired at a resolution of 60,000 at m/z 200. Up to 15 of the most abundant isotope patterns with charges 2-5 from the survey scan were selected with an isolation window of 0.7 m/z and fragmented by higher-energy collision dissociation with normalized collision energies of 32, while the dynamic exclusion was set to 35 s. The maximum ion injection times for the survey scan and dual MS (MS/MS) scans (acquired with a resolution of 30,000 at m/z 200) were 50 and 150 ms, respectively. The ion target value for MS was set to 3e6 and for MS/MS was set to 1e5, and the minimum AGC target was set to 1e3. The data were processed with MaxQuant v. 2.6.7.0 and the peptides were identified from the MS/MS spectra searched against the reviewed UniProt mouse proteome using the built-in Andromeda search engine. Reporter ion MS2-based quantification was applied with reporter mass tolerance = 0.003 Da and min. reporter PIF = 0.75. Cysteine carbamidomethylation was set as a fixed modification and methionine oxidation, asparagine/glutamine deamidation, and protein N-terminal acetylation were set as variable modifications. For in silico digests of the reference proteome, cleavages of arginine or lysine followed by any amino acid were allowed (trypsin/P), and up to two missed cleavages were allowed. The FDR was set to 0.01 for peptides, proteins, and sites. Since two raw files were collected (double injection of the sample), match between runs was enabled. Other parameters were used as preset in the software. Reporter intensity corrected values for protein groups were loaded into Perseus v. 1.6.10. Standard filtering steps were applied to clean up the dataset: reverse (matched to decoy database), only identified by site, and potential contaminant (from a list of commonly occurring contaminants included in MaxQuant) protein groups were removed. Reporter intensity corrected values were log2 transformed, and protein groups with values across all samples were kept. To determine which proteins were bound to Probe 1 vs spermidine, Student's t-test (1-sided, permutation-based FDR = 0.01, S0 = 0.5, n = 3) was performed. The table was exported from Perseus and formatted to its final form (Table S1) in Microsoft Excel 2016. The corresponding Volcano plot was created in Microsoft Excel 2016.

<u>Sample processing for proteomics after treatment in cells (peptide-level enrichment of biotinylated peptides)</u>

After ligation with azido-$PEG_3$-biotin and protein precipitation from HEK293T lysates, the pellets were resuspended in HEPES (100 mM, pH 8.0) containing TCEP (5 mM) and chloroacetamide (10 mM). Sequencing Grade Modified Trypsin (Promega) was added, and the sample was incubated overnight at 37 °C with 1100 RPM shaking. EDTA-free Protease Inhibitor Cocktail (1x, Roche) was added to quench residual trypsin activity. NeutrAvidin agarose resin was washed with 100 mM HEPES (3×). The tryptic peptide sample was centrifuged and the clear supernatant was added to the beads. The enrichment was carried out for 2 h at RT with 1100 RPM shaking. The supernatant was removed and the beads were washed with 100 mM HEPES (3×), with water (2×), and with 10% acetonitrile in water (2×). Probe-modified peptides were eluted at 37 °C with 0.1% trifluoroacetic acid and 0.2% formic acid in water (2×15 min). Combined eluates were concentrated using a SpeedVac concentrator. The sample was pre-cleaned using C18 StageTips and resuspended in 0.1% TFA and 2% acetonitrile in water before the LC-MS/MS measurement. Chromatographic separation was performed on an Easy-Spray Acclaim PepMap column (15 cm length × 75 µm inner diameter, Thermo Fisher Scientific) at 60 °C by applying 75 min acetonitrile gradients in 0.1% aqueous formic acid at a flow rate of 450 nl/min. An UltiMate 3000 nano-LC system was coupled to a Q Exactive HF-X mass spectrometer via an easy-spray source (all Thermo Fisher Scientific). The Q Exactive HF-X was operated in a data-dependent acquisition mode with survey scans acquired at a resolution of 60,000 at m/z 200. Up to 15 of the most abundant isotope patterns with charges 3-6 from the survey scan were selected with an isolation window of 1.3 m/z and fragmented by higher-energy collision dissociation with normalized collision energies of 27, while the dynamic exclusion was set to 35 s. The maximum ion injection times for the survey scan and MS/MS scans (acquired with a resolution of 15,000 at m/z 200) were 45 and 150 ms, respectively. The ion target value for MS was set to 3e6 and for MS/MS was set to 1e5, and the minimum AGC target was set to 1e3. The data were processed with MaxQuant v. 2.6.7.0 and the peptides identified from the MS/MS spectra were searched against reference Uniprot Human Proteome using the built-in Andromeda search engine. Cysteine carbamidomethylation, the cysteine-Probe1 thiotriazole adduct (Δmass = 625.3734), and Probe 1-derived hypusine (Δmass = 569.2996) or deoxyhypusine (Δmass = 553.3046) carrying the biotinylated triazole tag were set as variable modifications. The maximal number of variable modifications per peptide was set to 2. For in silico digests of the reference proteome, cleavages of arginine or lysine followed by any amino acid were allowed (trypsin/P), and up to two missed cleavages were allowed. The FDR was set to 0.01 for peptides, proteins, and sites. Other parameters were used as pre-set in the software. The modificationSpecificPeptides file (output from MaxQuant) was loaded into Perseus v. 1.6.10. Standard filtering steps were applied to clean up the dataset: reverse (matched to decoy database) and potential contaminant (from a list of commonly occurring contaminants included in MaxQuant) protein groups were removed. Data for peptides carrying the cysteine-Probe 1 thiotriazole adduct were exported from Perseus and formatted in Microsoft Excel 2016 to generate Table S2. Peptides modified by Probe 1-derived deoxyhypusine or hypusine carrying the biotinylated triazole tag were not detected.

<u>LC-MS/MS analysis of human recombinant eIF5A after in vitro reaction with DHS/DOHH</u>

eIF5A (20 µg) was incubated for 1 h at room temperature in glycine/NaOH buffer (0.2 M) containing NAD (1 mM) and either SPD (1 mM) or Probe 1 (1 mM), with or without DHS (2 µg) and DOHH (2 µg). The total reaction volume was 50 µl. The samples were diluted (1:4) with HEPES (100 mM, pH 8.0) and subjected to digestion with trypsin (sequencing grade, Promega) in the presence of TCEP (5 mM) and CAA (10 mM) overnight at 37 °C. The digested samples were desalted using C18 StageTips, concentrated, and resuspended in 0.1% TFA and 2% acetonitrile in water prior to LC-MS/MS analysis. Chromatographic separation was performed on an Easy-Spray Acclaim PepMap column (50 cm length × 75 µm inner diameter, Thermo Fisher Scientific) at 55 °C by applying 90 min acetonitrile gradients in 0.1% aqueous formic acid at a flow rate of 300 nl/min. An UltiMate 3000 nano-LC system was coupled to a Q Exactive HF-X mass spectrometer via an easy-spray source (all Thermo Fisher Scientific). The Q Exactive HF-X was operated in a data-dependent acquisition mode with survey scans acquired at a resolution of 60,000 at m/z 200. Up to 15 of the most abundant isotope patterns with charges 2-6 from the survey scan were selected with an isolation window of 1.3 m/z and fragmented by higher-energy collision dissociation with normalized collision energies of 27, while the dynamic exclusion was set to 20-35 s. The maximum ion injection times for the survey scan and MS/MS scans (acquired with a resolution of 15,000 at m/z 200) were 45 and 96 ms, respectively. The ion target value for MS was set to 3e6 and for MS/MS was set to 1e5, and the minimum AGC target was set to 1e3. The data were processed in MaxQuant v. 2.6.7.0, and peptides were identified from MS/MS spectra searched against the UniProt *E. coli* reference proteome (human proteins were expressed in *E. coli*) supplemented with the sequences of human DOHH, DHS, and EIF5A using the built-in Andromeda search engine. Cysteine carbamidomethylation was set as a fixed modification, whereas hypusine (Δmass = 87.0684), deoxyhypusine (Δmass = 71.0735), and their N-propargylated forms (Δmass = 125.0841 and Δmass = 109.0892), when Probe 1 was used, were set as variable modifications. For in silico digestion of the reference proteome, cleavages C-terminal to arginine or lysine followed by any amino acid were allowed (trypsin/P), with up to three missed cleavages. The FDR was set to 0.01 at the peptide, protein, and site levels. All other parameters were kept at their default values. The modificationSpecificPeptides output from MaxQuant was imported into Perseus v. 1.6.10. Peptides with gene names matching EIF5A were retained, and intensity values were $\log_2$-transformed. For unmodified peptide analysis, peptides that did not cover the known or putative modification site were required to have values across all compared conditions. For peptides covering the known or putative modification site, in the rare cases in which the value in the complete reaction sample (containing DHS and DOHH during incubation) was missing, intensities were imputed from a normal distribution (downshift = 1.8 SD, width = 0.3 SD). Median intensity values were subtracted within each sample, and the resulting data were exported to Microsoft Excel 2016 for final formatting (Table 3). MS/MS spectra were generated in MaxQuant Visualization.
